# A Method for Cost-Effective and Rapid Characterization of Engineered T7-based Transcription Factors by Cell-Free Protein Synthesis Reveals Insights into the Regulation of T7 RNA Polymerase-Driven Expression

**DOI:** 10.1101/614545

**Authors:** John B. McManus, Richard M. Murray, Peter A. Emanuel, Matthew W. Lux

**Affiliations:** US Army Combat Capabilities Development Command, Army Research Laboratory, 2800 Powder Mill Rd, Adelphi, MD 20783; California Institute of Technology, Biology and Biological Engineering, 1200 East California Blvd, Pasadena, CA 91125; US Army Combat Capabilities Development Command Chemical Biological Center, 8567 Ricketts Point Road, APG, MD 21010

**Keywords:** cell-free protein synthesis, TXTL, T7 RNA Polymerase, tetracycline

## Abstract

The T7 bacteriophage RNA polymerase (T7 RNAP) serves as a model for understanding RNA synthesis, as a tool for protein expression, and as an actuator for synthetic gene circuit design in bacterial cells and cell-free extract. T7 RNAP is an attractive tool for orthogonal protein expression in bacteria owing to its compact single subunit structure and orthogonal promoter specificity. Understanding the mechanisms underlying T7 RNAP regulation is important to the design of engineered T7-based transcription factors, which can be used in gene circuit design. To explore regulatory mechanisms for T7 RNAP-driven expression, we developed a rapid and cost-effective method to characterize engineered T7-based transcription factors using cell-free protein synthesis and an acoustic liquid handler. Using this method, we investigated the effects of the tetracycline operator’s proximity to the T7 promoter on the regulation of T7 RNAP-driven expression. Our results reveal a mechanism for regulation that functions by interfering with the transition of T7 RNAP from initiation to elongation and validates the use of the method described here to engineer future T7-based transcription factors.

**Highlights:**

- Development of a rapid and cost-effective method for screening synthetic promoters.
- Insights into the regulation of engineered T7-based transcription factors and T7 RNAP enzyme kinetics.
- Validation of this method by comparison with the T7 RNAP kinetic model.

## Introduction

Since its isolation in 1970 [1], the T7 bacteriophage RNA polymerase (T7 RNAP) has become a model for understanding RNA synthesis, as well as emerging as an important tool for protein expression [2] and as an actuator for synthetic gene circuit design in bacterial cells and cell-free extract [3,4]. The T7 RNAP is a 98 kDa, single-subunit enzyme that requires no additional protein factors to perform the complete transcriptional cycle [5]. This transcriptional cycle can be broken into three phases: binding, initiation, and elongation. During binding, T7 RNAP specifically recognizes the T7 promoter [5], and it has little affinity for other sequences, even the closely-related T3 promoter [6]. T7 RNAP then performs several rounds of abortive transcription, producing transcripts 10-12 nucleotides in length [7,8]. The enzyme undergoes a conformational change that marks its transition from initiation to the highly-processive elongation phase [9], transcribing RNA at a rate of 43 nucleotides per second [10], and producing transcripts greater than 10 kb in size [11]. Comparing these characteristics with the native *Escherichia coli* RNAP, which requires multiple subunits and recognizes orthogonal promoters, demonstrates the utility of T7 RNAP as an orthogonal tool. T7 RNAP was exploited early and often in synthetic gene circuit design owing to its ability to partially insulate circuit function from the host metabolism [3,4,12–14]. On average, there are only 2,000 native RNAP molecules per *E. coli* cell [15], and thus, the fluctuation of intracellular resources and variation in drag on metabolism, corresponding with growth phase and conditions, can complicate the predicted function of a gene circuit. Having an orthogonal tool set of well-characterized actuators, such as T7 RNAP, is vital to the ability to accurately predict gene circuit function [12].

One major drawback of the T7 RNAP system is the lack of regulatory mechanisms, beyond regulating the expression level of T7 RNAP, and a small set of synthetic promoters [4]. This stands in stark contrast to the large libraries of available and regulatable native promoters, including sets that are highly characterized and lack crosstalk [16]. Understanding regulatory mechanisms for T7 RNAP is important to the discovery of new engineered T7-based transcription factors that can be used in gene circuit design. Here we developed a rapid and cost-effective method to characterize promoter-operator combinations using cell-free protein synthesis and an acoustic liquid handler. We chose, as a model to test this method, the tetracycline regulatory (*tet*) system. The *tet* system functions to regulate protein expression when the homodimeric tetracycline repressor protein (TetR) binds to the tetracycline operator (*tetO*) sequence, resulting in downregulated transcription [17,18]. The *tet* system has been exploited in synthetic gene circuit design to regulate both host RNAP and T7 RNAP-driven expression [4,17,19–22]. In the case of host RNAP regulation, the *tetO* sequence can be placed between the −10 and −35 regions, resulting in relatively greater dynamic ranges when compared with T7 RNAP regulation by the same system [23]. Due to the single binding region of the T7 promoter, the *tetO* sequence can only be placed up or downstream from the T7 promoter. Thus, we investigated the effects of proximity of the *tetO* sequence to the T7 promoter on the regulation of T7 RNAP-driven expression.

We show that, irrespective of the position of *tetO* within the first 13 bp downstream of the T7 promoter, T7 RNAP-driven expression is downregulated equally by TetR. Conversely, placing the *tetO* sequence upstream from the T7 promoter shows nearly equal expression in the presence or absence of TetR. Our results suggest that *tet* regulation of T7 RNAP occurs by interfering with the initiation phase of T7 RNAP. We believe that this finding reveals characteristics regarding regulation of T7 RNAP that can be used in the engineering of T7-based transcription factors, and that such engineered transcription factors can be rapidly characterized by the methods described herein.

## Materials and Methods

### PCR-Amplification of Linear Template

Linear template for use in cell-free protein synthesis was amplified in two rounds of PCR using a single universal reverse primer and a set of two overlapping forward primers (one for each round of PCR) containing different positional pairings of T7-*tetO* sequences. PCR products were isolated by the Qiagen gel purification kit (Qiagen), quantified using a nanodrop spectrophotometer, diluted to 20 nM, and stored at −20°C until use (Fig. S2).

### Cloning of Linear Constructs into PY71

Linear templates for each *tetO* position were PCR-amplified using forward and reverse primers to add homology regions to the 5’ and 3’ ends in order to facilitate seamless cloning into the pY71 vector (Fig. S1). Linear templates were cloned into the pY71 vector using the In-fusion seamless cloning kit (Clontech). Circular templates were amplified in *E. coli* DH5α cells and isolated by Qiagen miniprep kit (Qiagen), quantified using a nanodrop spectrophotometer, diluted to 20 nM, and stored at −20°C until use.

### Preparation of Cell-Free Extract and Cell-Free Reactions

Cell-free extract was prepared according to Sun et al. [24] with the modification that cells were lysed by French pressure cell at 10,000 psi rather than by bead beating. Cell-free reactions were prepared according to Sun et al. [24]. Except where noted, a reaction contained energy buffer (3.3 µL), extract (4.2 µL), T7 polymerase (0.12 µL from 13 mg/mL stock), malachite green (0.2 µL from 10 mM stock), GamS (0.15 µL from 207 µM stock), DNA template (1 µL), and water (0.03 µL). Reactions were distributed by electronic pipette and TetR dilutions were distributed into cell-free reactions in volumes of 1 µL using an Echo 525 acoustic liquid handler (Labcyte Inc.).

sfGFP expression, in a total volume of 10 µL, was measured in black, clear-bottom 384 well plates (Greiner). Reactions were performed at 30°C and terminated after 12 h. sfGFP expression was measured by fluorescence in a Biotek H1 plate reader at 415 nm (ex) / 528 nm (em). Where indicated, RFU values were converted to protein concentration using calibration curves generated with purified sfGFP (a gift from Scott Walper, Naval Research Lab).

### Purification of TetR Protein

The *tetR* gene was PCR amplified from *E. coli* DH5α total DNA and seamlessly cloned into a pET-22B vector containing a C-terminal hexahistidine tag and expressed in *E. coli* BL21(DE3) cells. Cells (1 g) were resuspended in lysis buffer (5 mL) (50 mM Tris-Cl, 500 mM NaCl, 5 mM imidazole, pH 8.0) and lysed by sonication. The lysate was clarified by centrifugation at 15,000 × g for 30 minutes at 4°C. The supernatant (5 mL) was mixed with Ni-NTA resin (1 mL) (Sigma Aldrich), incubated at 4°C for 1 h, and loaded into a column. The resin was washed with 10 column volumes of wash buffer (50 mM Tris-Cl, 500 mM NaCl, 25 mM imidazole, pH 8.0). TetR was collected with three column volumes of elution buffer (50 mM Tris-Cl, 500 mM NaCl, 250 mM imidazole, pH 8.0) and concentrated to 1.5 mL using a 3 kDa cut-off centrifugal concentrator (Millipore). TetR was dialyzed against 2 L of dialysis buffer (50 mM NaHPO_4_, 100 mM NaCl, 2% DMSO, pH 7.5) for 1 h at 4°C then 2 L of dialysis buffer overnight at 4°C. TetR was then centrifuged at 14,000 × g for 10 minutes to remove precipitate and quantified by absorbance at 280 nm using the molar extinction coefficient (15,845 M^-1^ cm^-1^). Working TetR dilutions were first prepared, using dialysis buffer, by serial dilution, then flash frozen to reduce error and stored at −80°C until use.

### Purification of GamS Protein

The GamS protein was expressed from the pBad vector according to Sun et al. [25] and purified by nickel affinity as described for the TetR protein. GamS was dialyzed against 2 L of dialysis buffer (50 mM NaHPO_4_, 1 mM DTT, 1 mM EDTA, 100 mM NaCl, 2% DMSO, pH 7.5) for 1 h at 4°C then 2 L of dialysis buffer overnight at 4°C. GamS was then centrifuged at 14,000 × g for 10 minutes to remove precipitate and quantified by absorbance at 280 nM using the molar extinction coefficient (11,460 M^-1^ cm^-1^). GamS was diluted in dialysis buffer to 207 µM, flash frozen in liquid nitrogen, and stored at −80°C until use.

### Purification of T7 RNA Polymerase

T7 RNAP was expressed and purified according to Swartz et al. [26]. T7 RNAP was diluted to 13 mg/mL, flash frozen in liquid nitrogen, and stored at −80°C until use.

### Curve Fitting and Statistical Analysis

sfGFP expression curves were fit to a sigmoidal equation:

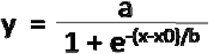

Four parameter logistic regressions were performed using Prism software (Graphpad). The maximum repression value for each construct was expressed as the difference between the top and bottom values determined by the four parameter logistic fit.

Statistical analysis was performed by a one-way ANOVA test, a two-way ANOVA test, or by Welch’s t-test, as indicated in the results and figures, using Prism software (Graphpad). For ANOVA tests, Tukey’s method was applied to determine statistical significance between data at each *tetO* position.

## Results

### TetO Represses T7 RNAP Equally when It is Positioned within the First 13 Bases of the T7 Transcript

In order to effectively explore and understand the design space of engineered T7-based transcription factors, we aimed to develop a method that would allow for rapid and cost-effective characterization of promoter-operator combinations. To do this, we generated linear template by PCR-amplifying the sfGFP gene from the sfGFP-containing pY71-GFP plasmid using one universal reverse primer and different forward primers containing spatial combinations of T7-*tetO* across a stretch of standard base pairs (Fig. S2 and S3). We chose the *tetO*_*2*_ sequence for this work because TetR has shown approximately twice the affinity for *tetO*_*2*_ over *tetO*_*1*_ [27]. We chose cell⍰free extract, which is prepared from *E. coli* cells, in-house, as a rapid and cost-effective medium for the measurement of protein expression [24,25]. This method circumvents the need to clone, transform, and measure expression in whole cells, allowing for the characterization of many promoter-operator combinations in a single day. Additionally, it allowed us to directly probe the effects of the *tetO* position using purified TetR. For nomenclature purposes, constructs were designated according to the number of base pairs that the 5’ end of the *tetO* sequence lies away from the transcriptional start site of the T7 promoter sequence (or the length of the transcript, in bases) (Fig. S3).

In order to prevent the degradation of linear template, GamS protein was added to the reaction mixture [25]. Different concentrations of TetR were added to reaction wells using an acoustic liquid handler. Cell-free reactions were run for 12 h, expression curves were fit using a sigmoidal regression, and the maximum sfGFP expression values were used for further evaluation (Fig. 1A).

**Figure 1.**
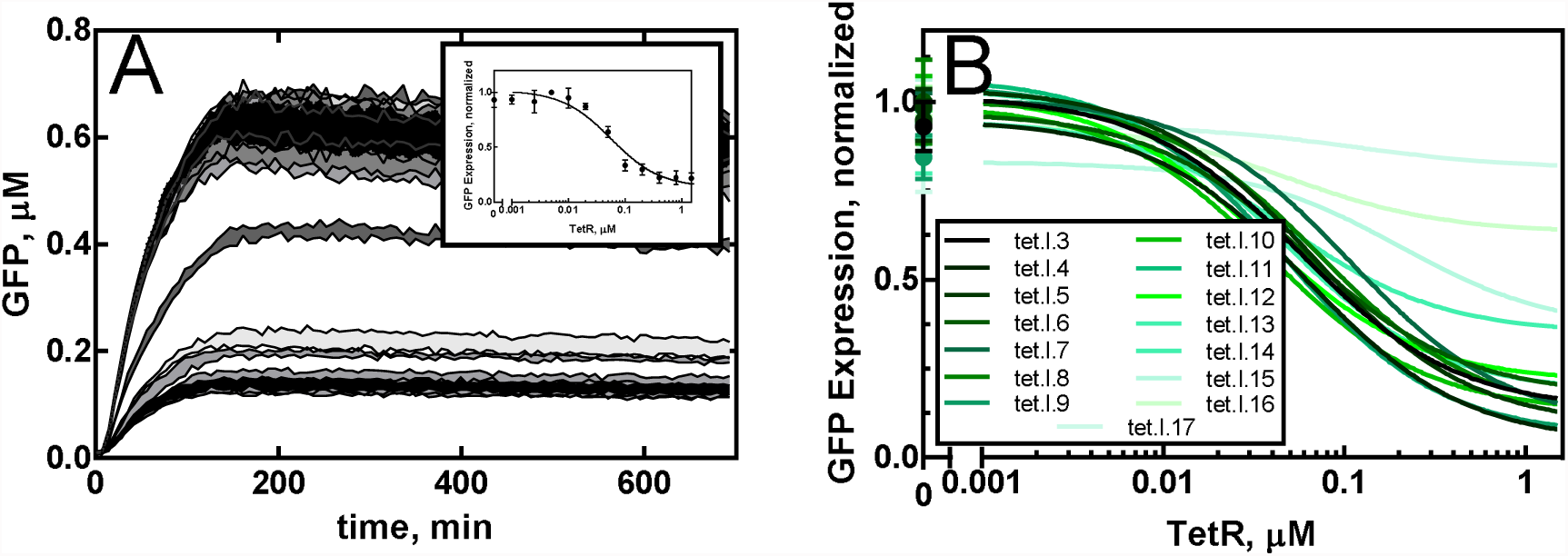
GFP Expression and TetR Dose-Response from Linear Template. **(A)** GFP was expressed from 2 nM linear tet.I.3 template in 10 µL of cell-free extract containing varying concentrations of TetR. Each trace represents the mean and standard deviation of three reactions. **(A, inset)** The expression curves shown in panel A were fit to a sigmoid regression. The maximum expression values were rescaled and plotted against TetR concentration and a four parameter logistic curve fit was applied. **(B)** The same analysis was performed for each *tetO* position and only the fits are shown.

Despite that the same concentration of template was added to each reaction, we observed variation in the maximum sfGFP expression using different templates, ranging from 0.4 µM to 1.4 µM, in the absence of TetR (Fig. S4A). To determine whether these fluctuations were simply due to the positioning of the *tetO* sequence, we tested two different preparations of each template with no TetR present (Fig. S4A). When a one-way ANOVA test was applied to the expression data from lot 1, no apparent pattern that might indicate an effect of the *tetO* position on expression became apparent (Fig. S4C). A two-way ANOVA test was applied to the expression data of both lots in order to determine statistical significances in the difference in expression at each *tetO* position. A pattern indicating no significance, running along the diagonal in Figure S4E, would illustrate the reproducibility of the variation in expression between lots (Fig. S4A). However, such a pattern does not emerge, suggesting that variation cannot be explained by the *tetO* position alone.

In order to facilitate direct comparisons between each *tetO* position, all mean expression values were divided by the greatest mean expression value for that position. This transformation rescaled the expression data to a maximum value of one for each position. Rescaled expression values were then plotted against the TetR concentration to generate dose-response profiles for each *tetO* position (Fig. 1A, inset). Each dose-response profile was fit to a four parameter logistic regression curve (Fig. 1B) in order to obtain maximum repression values and 1/2 inhibitory concentration (IC_50_) values for TetR for each construct (Table S1).

A one-way ANOVA test comparing IC_50_ values at each position, revealed no significant difference (P ≤ 0.0001), with the exception being that of tet.I.17, comparatively to other *tetO* positions (Fig. 2A). However, this may be explained by the difficulty in fitting the dose-response profile for tet.I.17, resulting in a large standard deviation (Fig. 2A). These results suggest that there is no effect of the *tetO* position on TetR binding, using linear template.

**Figure 2.**
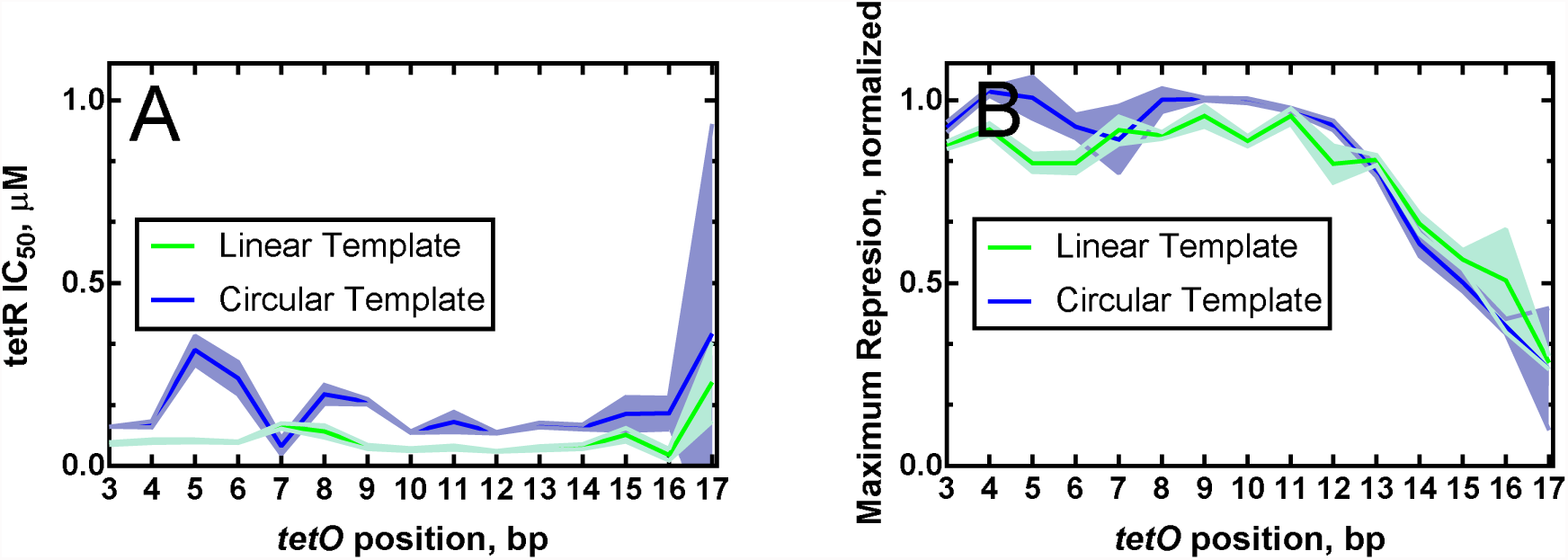
The Effect of *tetO* Position and Template Type on T7-Driven Expression in Cell-Free Extract. **(A)** The TetR IC_50_ values for each construct, calculated from the four parameter logistic curve fits (Fig. 1 and 3) were plotted against *tetO* position. Each trace represents the mean and standard deviation of three replicates. **(B)** The maximum repression values for each construct, calculated by subtracting the minimum value from the maximum value for each logistic curve fit (Fig. 1B and 3B) were plotted against *tetO* position. Each trace represents the mean and standard deviation of three replicates.

When a one-way ANOVA test to compare maximum repression values at each *tetO* position was applied, only tet.I.15 through tet.I.17 showed a significant difference (P ≤ 0.0001) on downregulation of T7 RNAP-driven expression (tet.I.14 falls at a transitional position (P ≤ 0.01)) (Fig. 2B). There is little, if any, observable effect of the *tetO* position on downregulation when the *tetO* sequence is less than 14 bp downstream from the T7 promoter. This phenomenon is best illustrated by the heat map in Figure 4A. Our results suggest one of two mechanisms for regulation may be at play: (1) that TetR blocks the binding of T7 RNAP, equally, up through position 13 or (2) that TetR prevents T7 polymerase from transitioning from initiation to elongation.

**Figure 3.**
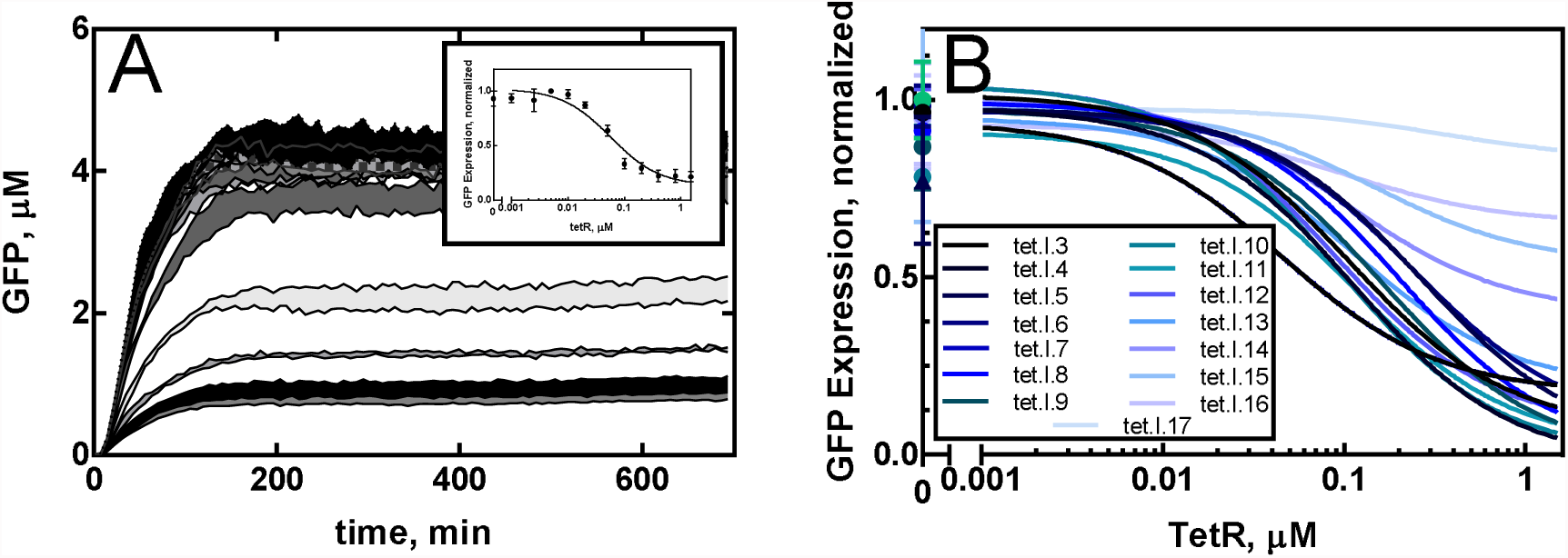
GFP Expression and TetR Dose-Response from Circular Template. **(A)** GFP was expressed from 2 nM circular PY71-tet.I.3 template in 10 µL of cell-free extract containing varying concentrations of TetR. Each trace represents the mean and standard deviation of three reactions. **(A, inset)** The expression curves shown in panel A were fit to a sigmoid regression. The maximum expression values were rescaled and plotted against TetR concentration and a four parameter logistic curve fit was applied**. (B)** The same analysis was performed for each *tetO* position and only the fits are shown.

**Figure 4.**
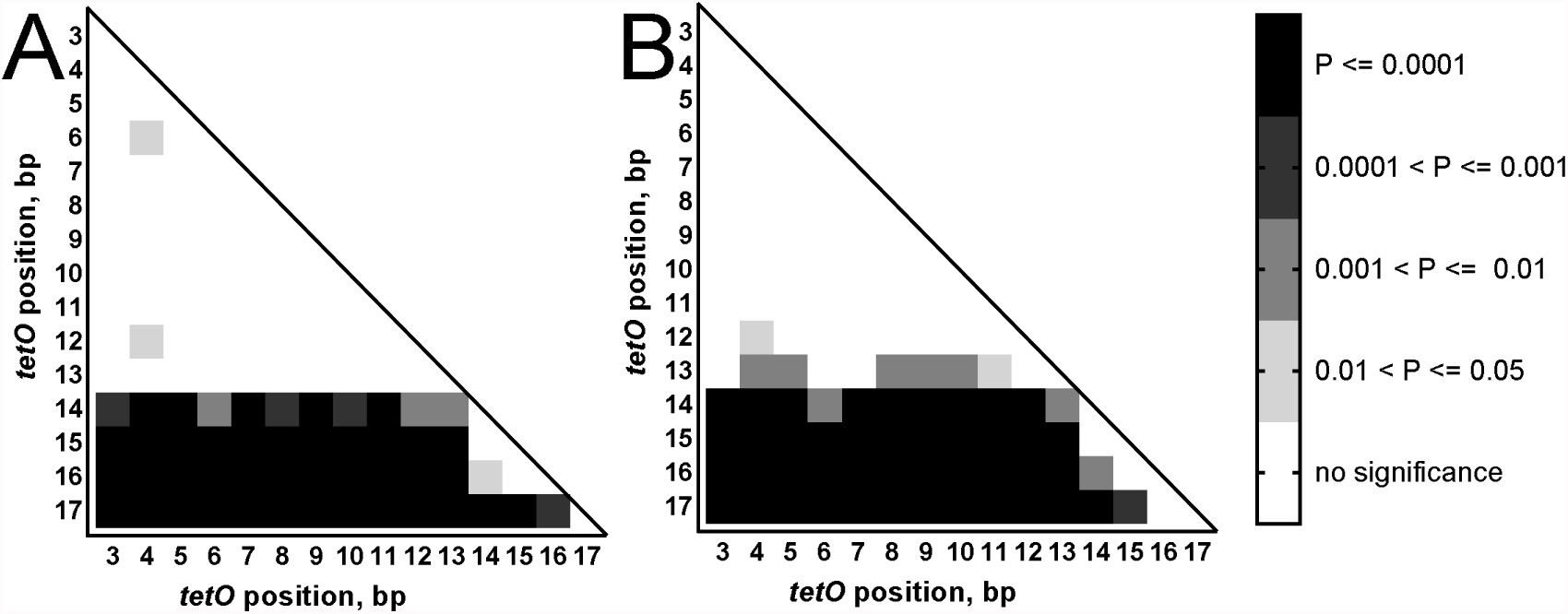
Statistical Analysis of tetO Positioning Effects on T7-Driven Expression. The maximum repression values calculated for both **(A)** linear and **(B)** circular templates were subject to an ordinary ANOVA analysis. Adjusted P-values (alpha = 0.05) were converted to a heat map and plotted comparatively against each construct to demonstrate significance.

### Template Format Impacts TetR Binding to the TetO Sequence but Not Downregulation

To investigate the effects of template format (linear versus circular) on *tet* regulation of T7 RNAP-driven expression, each construct was seamlessly cloned into the pY71 expression vector, as described in Materials and Methods. Circular templates are differentiated from linear templates using the prefix pY71. Each circular template was evaluated for expression in cell-free extract, as described for linear templates. Different concentrations of TetR were added to reaction wells using an acoustic liquid handler. Cell-free reactions were run for 12 h, expression curves were fit using a sigmoidal regression, and the maximum sfGFP values were used for further evaluation (Fig. 3A)

We again observed variation in maximum sfGFP expression between templates, ranging from 0.5 µM to 4.4 µM in the absence of TetR. Again, we tested two different preparations of each template with no TetR present (Fig. S4B). When a one-way ANOVA test was applied to the expression data from lot 1, no pattern that might indicate an effect of *tetO* position on expression became apparent (Fig. S4D). A two-way ANOVA test was applied to the expression data of both lots in order to determine statistical significances in the difference in expression at each *tetO* position (Fig. S4F). In contrast to linear template, a pattern of no significance running along the diagonal in Figure S4F was observed (with the exception of pY71⍰tet.I.17). However, at most positions there is no statistically significant difference in expression. This may be attributed to the large standard deviations for the expression means of lot 2. Therefore, it is difficult to conclude that the variation in expression is necessarily due to *tetO* position alone, in circular template.

Expression values for circular template were rescaled as described for linear template. This was also useful for the direct comparison of data from circular template with those from linear template. Rescaled expression values were plotted against TetR concentration to generate dose-response profiles for each *tetO* position. Each dose-response profile was fit to a four parameter logistic regression curve (Fig. 3B) in order to obtain maximum repression values and IC_50_ values for TetR for each position (Table S2).

As with linear template, a one-way ANOVA test comparing IC_50_ values for all positions, in circular templates, revealed no significant difference (P ≤ 0.0001), except for the IC_50_ values for PY71-tet.I.5 and PY71-tet.I.6 comparatively to each other and the other *tetO* positions. Interestingly, a two-way ANOVA test, comparing IC_50_ values from circular and linear template, revealed a statistically significant difference. Further, with the exception of tet.I.15 and tet.I.17, a t-test of individual IC_50_ values between the two template formats, at each *tetO* position, shows a statistically significant difference (P ≤ 0.05). This suggests that, while *tetO* position does not influence TetR binding, template format may affect TetR binding. This is illustrated, qualitatively, by Figure 2A, which shows the relatively small variation in IC_50_ values along *tetO* position within template format compared with the relatively greater variation in IC_50_ values observed between template format.

Even though TetR displays an unusually low background affinity for DNA (∼10^5^ M^-1^) [28], we reasoned that some non-specific DNA binding might well account for the increased IC_50_ values in circular template, increasing the effective concentration required for downregulation. In order to test this hypothesis, we added 2 nM PCR-amplified linear pY71 backbone to 2 nM linear tet.I.5 template in cell-free reactions, while varying TetR concentration (Fig. S5). Our results showed that the IC_50_ value increased from 0.010 ± 0.002 µM to 0.029 ± 0.002 µM upon the addition of pY71 backbone DNA, a statistically significant increase (Welch’s t-test, p = 0.0006). The IC_50_ value for circular template in this experiment was 0.073 ± 0.047 µM, which is not statistically different from the IC_50_ value for linear template with linear pY71 backbone DNA (Welch’s t-test, p = 0.24). These results suggest that non-specific binding of TetR to the vector backbone may indeed be responsible for the observed differences between linear and circular templates. However, a t-test, applied to the IC_50_ values of tet.I.5 and pY71-tet.I.5 also shows no statistically significant difference (Welch’s t-test, p = 0.15). Thus, we cannot rule out contribution from other factors.

A one-way ANOVA test comparing TetR downregulation in circular template yielded results similar to that for linear template. This, again, is best illustrated by the heat map in Figure 4B, showing no statistically significant (P ≤ 0.0001) reduction in repression when *tetO* is upstream of position 14. A qualitative comparison of the heat maps in Figure 4 and the traces in Figure 2B show remarkably similar trends for both template formats. This suggests that template format has little, if any, impact on *tet* regulation of T7 RNAP-driven expression, verifying our use of linear template to evaluate engineered T7-based transcription factors.

### TetR Acts to Regulate T7-Driven Expression by Interfering with the Transition of T7 RNAP from Initiation to Elongation

In order to further investigate the *tet* regulatory mechanism for T7 RNAP, we placed the *tetO* sequence at positions 18, 27, and −36. For positions 18 and 27, we predicted that the trend of decreased regulation at positions further downstream from position 14 would continue. As expected, we observed nearly no downregulation at positions 18 and 27 (Fig. 5). Evaluation of position −36, which is immediately upstream of the T7 promoter sequence, was intended to differentiate the aforementioned competing hypotheses: (1) that the mechanism of regulation was either due to competitive binding between TetR and T7 RNAP near the promoter region, or (2) that TetR prevents T7 RNAP from transitioning from initiation to elongation. Interestingly, we found that, when the *tetO* sequence was placed immediately upstream from the T7 promoter sequence (tet.I.-36), TetR did not significantly downregulate T7 RNAP-driven expression (Fig. 5), suggesting that *tet* regulates T7 RNAP-driven expression by preventing T7 RNAP from transitioning from initiation to elongation, rather than blocking T7 RNAP binding. Our observations are supported by the findings of Iyer et al. [4] and are consistent with the kinetic model published by Skinner et al. [10].

**Figure 5.**
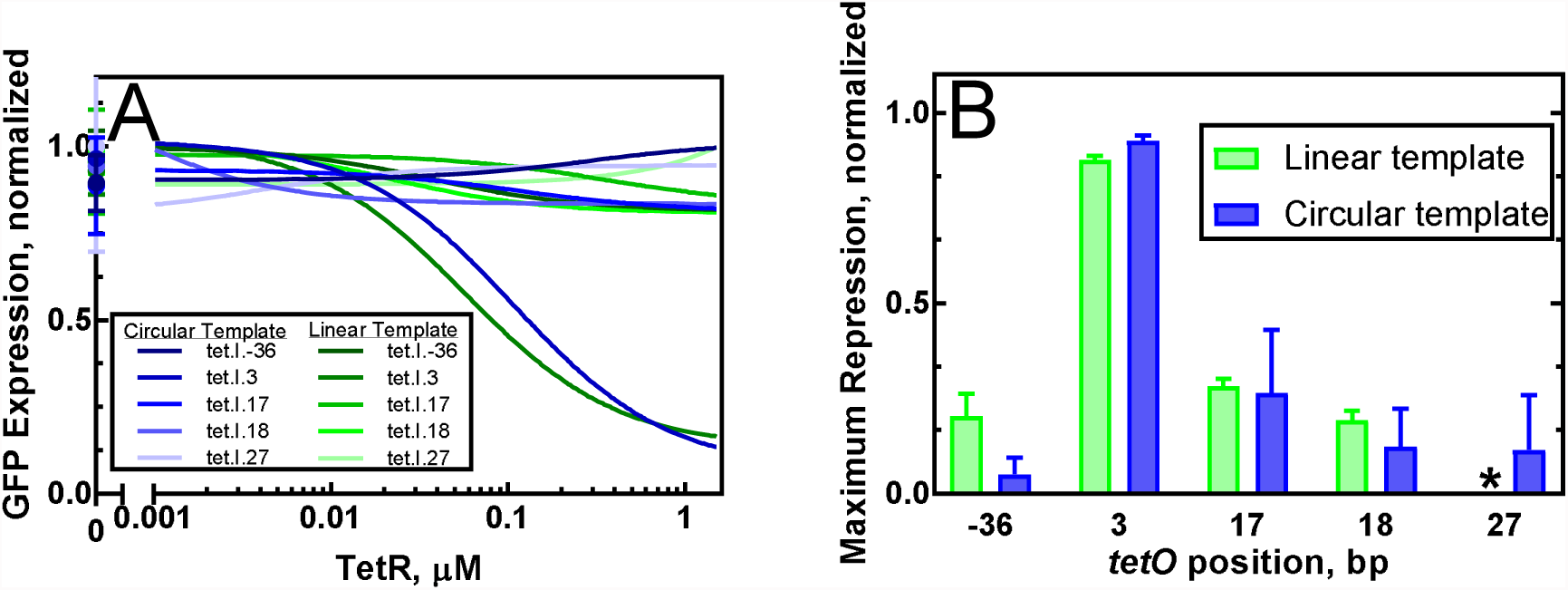
The Effect of Additional *tetO* Positions on T7-Driven Expression in Cell-Free Extract. **(A)** Expression curves for each template type were fit to a sigmoid regression. The maximum expression values were rescaled and plotted against TetR concentration and a four parameter logistic curve fit was applied. (B) The maximum repression values for each construct, calculated by subtracting the minimum value from the maximum value for each logistic curve fit in panel A, were plotted as bar graphs with the *tetO* position indicated on the x-axis. The template type is indicated in the legend. Each bar represents the mean and standard deviation of three replicates. *Logistic curve fit for tet.I.27 yielded a negative maximum repression.

## Discussion

Due to its orthogonality to bacterial host machinery, T7 RNAP is a powerful tool for gene circuit design, and regulating its activity is central to fine-tuning gene circuit function. As such, understanding regulatory mechanisms for T7 RNAP are important to the design of engineered T7-based transcription factors that can be used in synthetic gene circuits.

Here we describe a rapid and cost-effective method to characterize promoter-operator combinations using cell-free protein synthesis and an acoustic liquid handler. Using this method, we investigated the effect of proximity of the *tetO* sequence to the T7 promoter on the regulation of T7 RNAP-driven expression. We observed that the absolute expression levels of sfGFP varied between templates, even when the same amount of template was added. Therefore, we prepared a second lot of each template, and measured sfGFP expression in cell lysate. While it may be tempting to conclude that the *tetO* position is accountable for the variation in expression based on a qualitative assessment from the patterns in the traces of Figures S4E and S4F, statistical analyses of the expression data (Fig. S4) indicate that the *tetO* position is not alone responsible for the variation in expression. There are numerous factors, beyond minor variations in template sequence, that may be responsible for the observed variation. One such culprit is template preparation. Templates were prepared by the mini prep (Qiagen) method and small variations in the amount of salts carried over during template preparation may account, in part, for the variation in expression. It is known that the salts magnesium and potassium, which is contained as 0.9 M potassium acetate in the neutralization buffer of the Qiagen miniprep kit, are among the most important parameters, along with template concentration, that affect the efficiency of cell-free protein synthesis [24,29]. Despite variation in expression, when the expression values were rescaled, consistent patterns emerged that are useful in promoter characterization.

Our data comparing the IC_50_ values for TetR in both template formats raises the question of whether format influences the IC_50_ value and, by extension, TetR binding to the *tetO* sequence (Fig. 2A and S3). While our experiment of adding linear backbone DNA to linear template appears to at least partially explain the differences in IC_50_ values, a second explanation may be in play as well. The crystal structure of the TetR homodimer reveals binding to the *tetO* sequence via N-terminal alpha helices, which occupy the major grooves of the operator, engaging with all but three base pairs of the sequence [30]. In its relaxed state, the periodicity of the DNA helix is 10.4 bp per turn [31]. However, under supercoiled conditions, such as with circular DNA [32], the periodicity can vary between 10 to 11 bp turn [33], thus changing the width of the DNA grooves. It is plausible that the difference in IC_50_ values we observed between template formats can reasonably be attributed to the use of circular *versus* linear template. The contribution from experimental variation, however, makes it difficult to identify a single conclusive explanation.

In probing *tetO* position effects, we observed that T7 RNAP-driven expression is downregulated to the same degree when the *tetO* sequence is within 13 bp downstream from the T7 transcriptional start site (Fig. 3B) and that nearly no TetR downregulation is observed if the *tetO* sequenced is placed immediately upstream of the T7 promoter sequence (Fig. 5B). These results suggest that the *tet* regulatory mechanism for T7 RNAP operates by disrupting the transcriptional cycle, as described by Skinner et al. [10], at the initiation phase.

The transcriptional cycle, as described by Skinner et al. [10], proceeds through three phases: binding, initiation, and elongation (Fig. 6). During the binding phase, T7 RNAP recognizes the T7 promoter. Binding to the promoter sequence is close to the diffusion-controlled limit, indicating a relatively strong affinity of T7 RNAP for the T7 promoter [34] (Fig. 6A). Helix melting then occurs rapidly with the binding of the second ribonucleotide. Single molecule kinetic studies [10] on T7 RNAP revealed that, during initiation, T7 RNAP undergoes several rounds of abortive transcription across the first 12 bases of the template (Fig 6B), producing short RNA transcripts. Further, single molecule kinetics have shown that, during initiation, T7 RNAP favors dissociation (*k*_*off*_ = 2.9 s^-1^) over transitioning to elongation (*k*_*for*_ = 0.36 s^-1^) [10]. During initiation, T7 RNAP accommodates only three base pairs of the DNA-RNA heteroduplex within the active site of the enzyme [35]. This explains the relatively weak affinity of the T7 RNAP for the DNA template throughout the 12 bases that constitute initiation. As T7 RNAP transitions from initiation to elongation (Fig. 6C), it undergoes a conformational change: the collapse of the promoter binding site and the formation of a channel, around the active site, that accommodates seven base pairs of the DNA-RNA heteroduplex [9,36], as well as the formation of an N-terminal tunnel, allowing for the egress of the nascent RNA transcript [9]. Throughout elongation, processivity is increased significantly, indicating a relatively strong affinity of the T7 RNAP for the DNA template.

**Figure 6.**
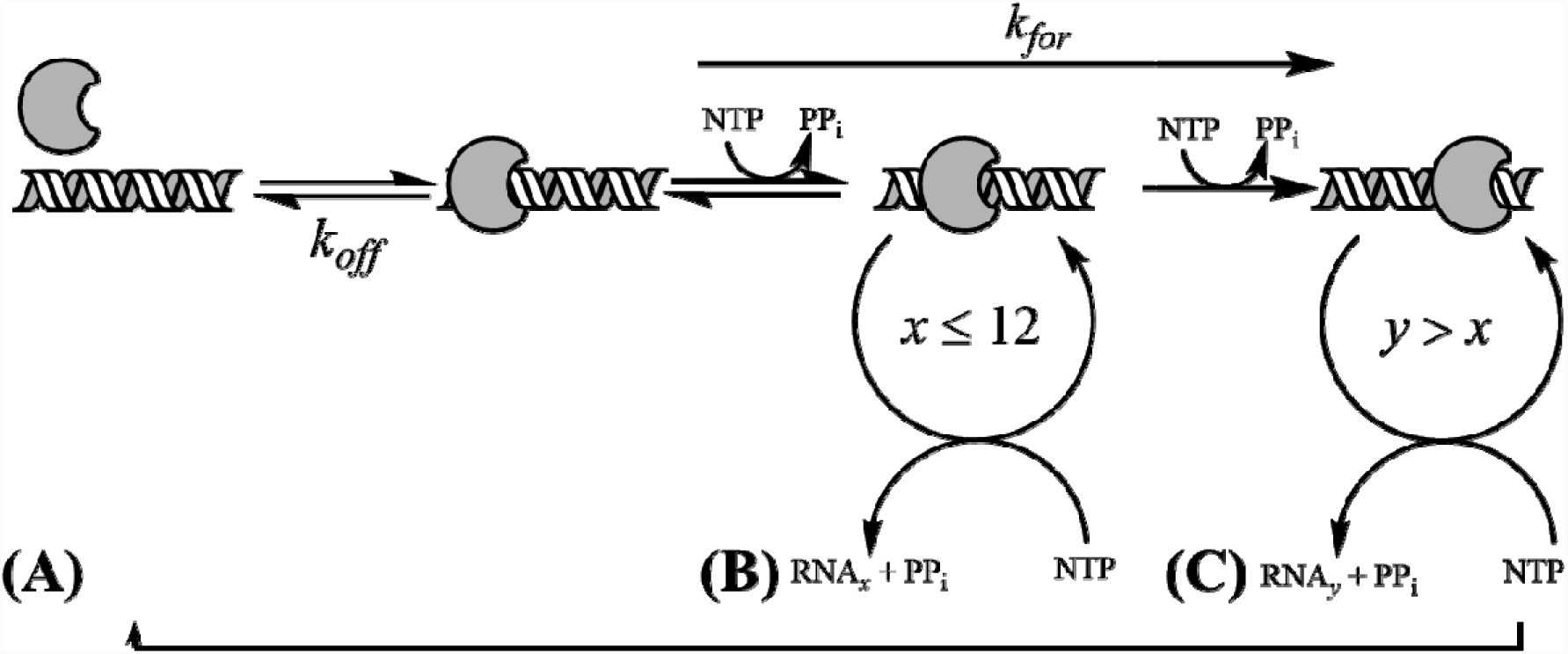
Kinetic Scheme for T7 RNAP-Catalyzed Transcription. **(A)** T7 RNAP binds to DNA template at the T7 promoter sequence. (B) T7 RNAP then undergoes initiation, during which short transcripts (RNA_x_) of no greater than 12 nucleotides (*x*) are synthesized. (C) After several rounds of abortive transcription, T7 RNAP enters elongation.

Our results are in good agreement with the transcriptional model presented by Skinner et al. [10], and illustrated in Figure 6. Consistent with the relatively weak affinity of T7 RNAP for the template during initiation, we observed that downregulation of T7 RNAP-driven expression is the strongest and identical, irrespective of the *tetO* position, as long as it is within the first 13 bp downstream from the T7 promoter (Fig. 3B and Fig. 4). TetR became less effective as *tetO* moved further downstream, consistent with T7 RNAP entering elongation following base 12 of its transcript [10], and the enzyme’s strong affinity for the DNA template during this phase. Finally, TetR was also not effective at downregulating expression when *tetO* was placed immediately upstream of the T7 promoter sequence (Fig. 5B), consistent with T7 RNAP’s strong affinity for the T7 promoter.

Understanding the mechanism of repression for T7 RNAP using well characterized systems, such as the *tet* system, will allow for the design of more effective engineered T7-based transcription factors. Our results suggest that the design of new repressor-based, T7-based transcription factors would be best narrowed to the initiation phase of T7 RNAP. Indeed, since *tet* repression is one of the most effective in native promoter systems [23], our results suggest that placing operator sites outside of the 12 bp stretch consistent with initiation is likely futile unless additional mechanisms such as DNA looping [4] are employed. We also showed that the method developed here, utilizing cell-free protein synthesis and linear template, can be used to rapidly evaluate any such new engineered T7-based transcription factors. These results will assist in expanding the pallet of engineered T7-based transcription factors for the design of gene circuits.

