## Supplementary material for "A Method for Cost-Effective and Rapid Characterization of Engineered T7-based Transcription Factors by Cell-Free Protein Synthesis Reveals Insights into the Regulation of T7 RNA Polymerase-Driven Expression": Supplimentary Material

TAGAAAAACTCATCGAGCATCAAATGAAACTGCAATTTATTCATATCAGGATTATCAATACCATATTTTTGAAAAAGCCGTTTCTGTAATGAAGGAGAAAACTCACCGAGGCAGTTCCATAGGATGGCAAGATCCTGGTATCGGTCTGCGATTCCGACTCGTCCAACATCAATACAACCTATTAATTTCCCCTCGTCAAAAATAAGGTTATCAAGTGAGAAATCACCATGAGTGACGACTGAATCCGGTGAGAATGGCAAAAGCTTATGCATTTCTTTCCAGACTTGTTCAACAGGCCAGCCATTACGCTCGTCATCAAAATCACTCGCATCAACCAAACCGTTATTCATTCGTGATTGCGCCTGAGCGAGACGAAATACGCGATCGCTGTTAAAAGGACAATTACAAACAGGAATCGAATGCAACCGGCGCAGGAACACTGCCAGCGCATCAACAATATTTTCACCTGAATCAGGATATTCTTCTAATACCTGGAATGCTGTTTTCCCGGGGATCGCAGTGGTGAGTAACCATGCATCATCAGGAGTACGGATAAAATGCTTGATGGTCGGAAGAGGCATAAATTCCGTCAGCCAGTTTAGTCTGACCATCTCATCTGTAACATCATTGGCAACGCTACCTTTGCCATGTTTCAGAAACAACTCTGGCGCATCGGGCTTCCCATACAATCGATAGATTGTCGCACCTGATTGCCCGACATTATCGCGAGCCCATTTATACCCATATAAATCAGCATCCATGTTGGAATTTAATCGCGGCTTCGAGCAAGACGTTTCCCGTTGAATATGGCTCATAACACCCCTTGTATTACTGTTTATGTAAGCAGACAGTTTTATTGTTCATGATGATATATTTTTATCTTGTGCAATGTAACATCAGAGATTTTGAGACACAACGTGGATCCTGCAGTTGAGATCCTTTTTTTCTGCGCGTAATCTGCTGCTTGCAAACAAAAAAACCACCGCTACCAGCGGTGGTTTGTTTGCCGGATCAAGAGCTACCAACTCTTTTTCCGAAGGTAACTGGCTTCAGCAGAGCGCAGATACCAAATACTGTCCTTCTAGTGTAGCCGTAGTTAGGCCACCACTTCAAGAACTCTGTAGCACCGCCTACATACCTCGCTCTGCTAATCCTGTTACCAGTGGCTGCTGCCAGTGGCGATAAGTCGTGTCTTACCGGGTTGGACTCAAGACGATAGTTACCGGATAAGGCGCAGCGGTCGGGCTGAACGGGGGGTTCGTGCACACAGCCCAGCTTGGAGCGAACGACCTACACCGAACTGAGATACCTACAGCGTGAGCATTGAGAAAGCGCCACGCTTCCCGAAGGGAGAAAGGCGGACAGGTATCCGGTAAGCGGCAGGGTCGGAACAGGAGAGCGCACGAGGGAGCTTCCAGGGGGAAACGCCTGGTATCTTTATAGTCCTGTCGGGTTTCGCCACCTCTGACTTGAGCGTCGATTTTTGTGATGCTCGTCAGGGGGGCGGAGCCTATGGAAACGAATTCAGATCTCGATCCCGCGA

**Figure S1. pY71 Sequence.**

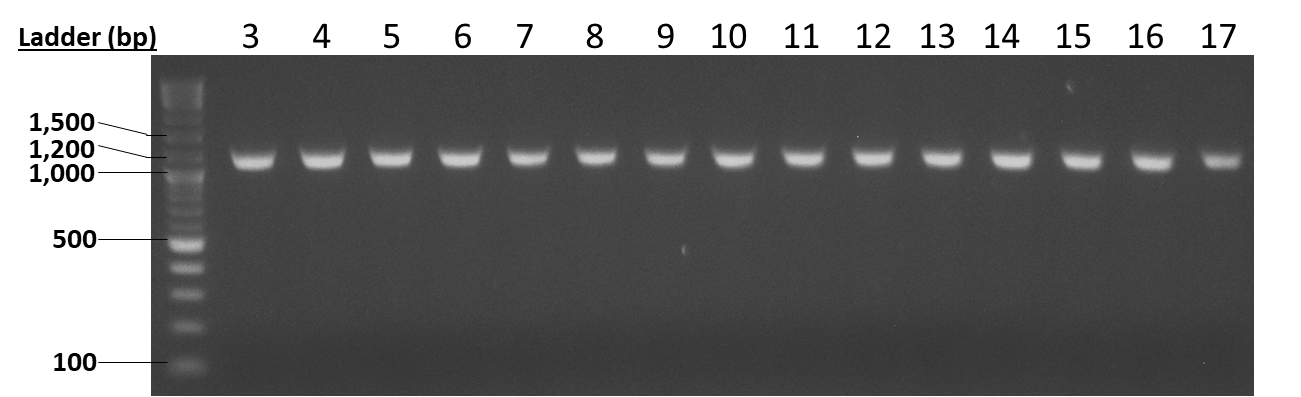

**Figure S2. Agarose Gel Electrophoresis of PCR-Amplified Template DNA.** Linear templates were PCR-amplified from circular template using identical reverse primers and forward primers containing T7-tetO combinations (the spacing between the two sequences is indicated by the number above each lane). Each amplicon was isolated using the Qiagen gel purification kit, as indicated in the Materials and Methods and diluted to 20 nM. Equal volumes of each template were loaded and separated on a 1% agarose gel. The calculated size for each amplicon is 1,126 bp.

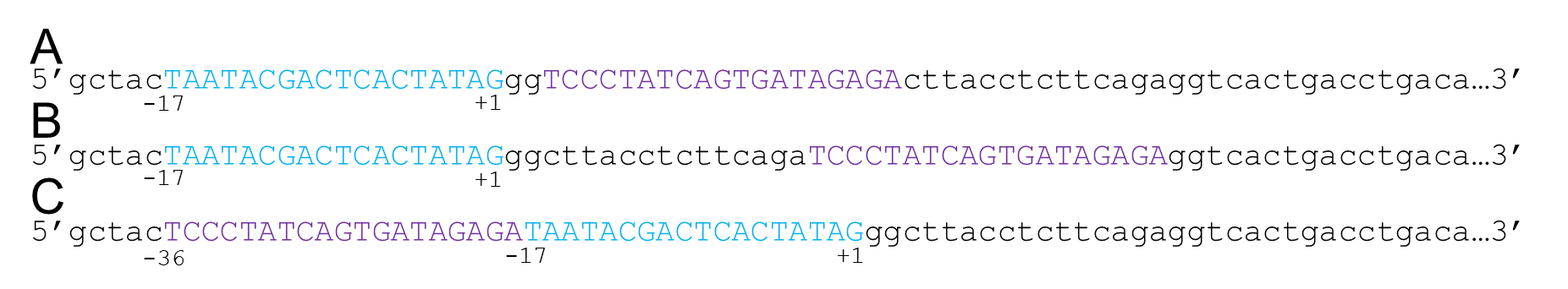

**Figure S3. Promoter Design.** Constructs were designated according to the number of base pairs that the 5’ end of the *tetO* sequence (purple) lies away from the T7 promoter (blue) transcriptional start site (+1). For example, **(A)** the construct containing *tetO* at the third base pair downstream from the T7 transcriptional start site was given the designation tet.I.3, while **(B)** *tetO* at the seventeenth base pair downstream from the T7 transcriptional start site was designated tet.I.17, and **(C)** the position of *tetO* immediately upstream of the T7 promoter sequence was designated tet.I.-36.

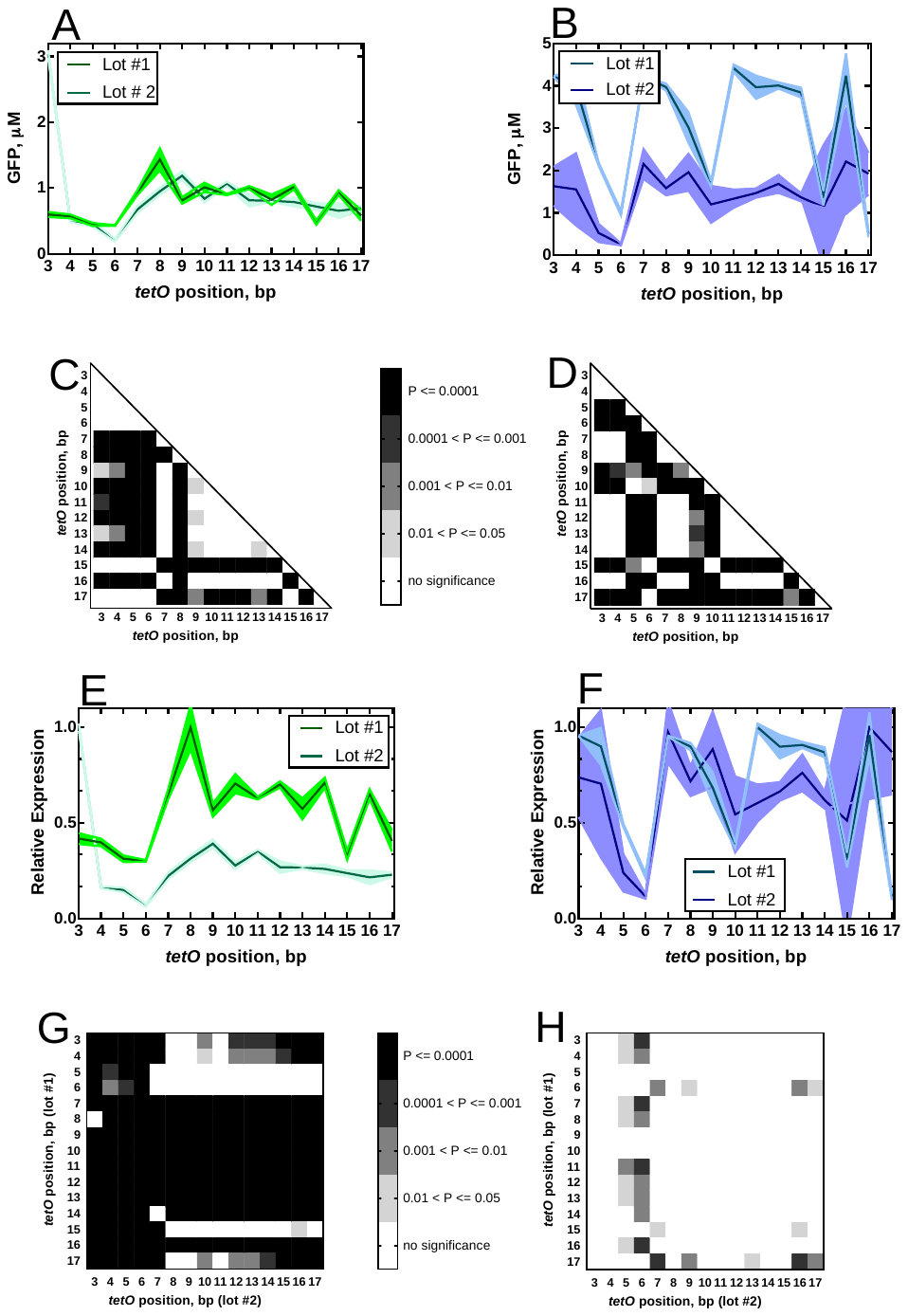

**Figure S4. An Analysis of Variation in Expression from Each Template.**  sfGFP was expressed from 2 nM **(A)** linear or **(B)** circular template and the expression curves were fit to a sigmoid regression. The maximum expression values from lot 1 or lot 2 template preparations were plotted against *tetO* position. The expression values at each *tetO* position for lots 1 in **(A)** and **(B)** were then compared by a one-way ANOVA analysis and adjusted P-values (alpha = 0.05) were converted to a heat map for both **(C)** linear and **(D)** circular templates. The maximum expression values from **(A)** and **(B)** were then rescaled, as described in Materials and Methods, to produce traces for **(E)** linear and **(F)** circular templates. Finally, the normalized expression values for lots 1 and 2 in **(E)** and **(F)** were subject to a two-way ANOVA analysis and adjusted P-values (alpha = 0.05) were converted to a heat map for both **(G)** linear and **(H)** circular templates. All traces represent the mean and standard deviation for three reactions.

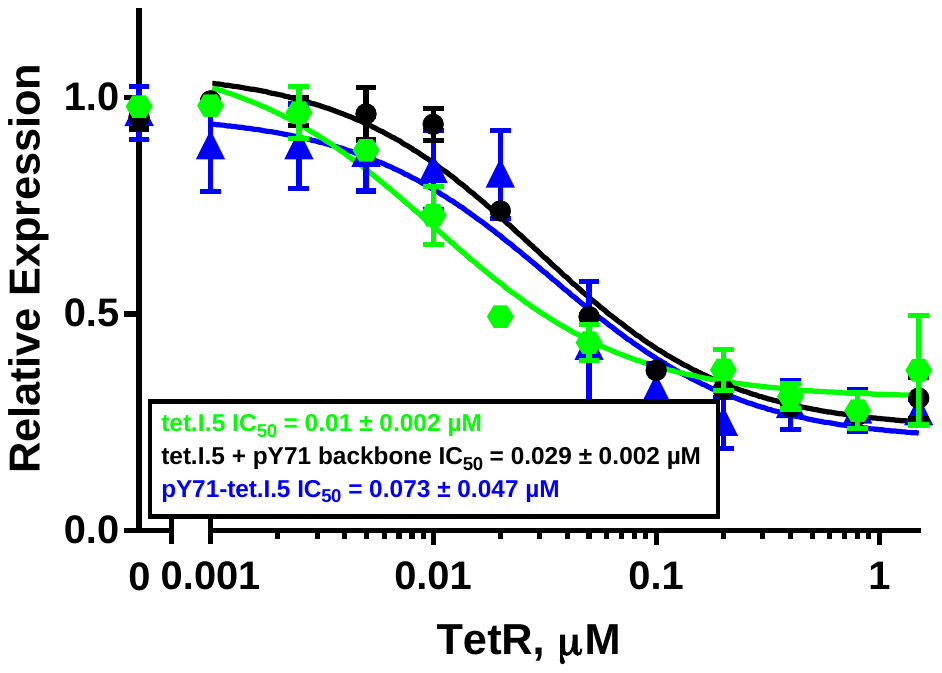

**Figure S5. TetR Dose-Response Curves for tet.I.5 (green hexagon), tet.I.5 + pY71 backbone DNA (black circles), and pY71-tet.I.5 (blue triangles).** GFP was expressed from 2 nM template in 10 µL of cell-free extract containing varying concentrations of tetR. Expression were fit to a sigmoid regression. The maximum expression values were rescaled and plotted against TetR concentration and a four parameter logistic curve fit was applied. Each point represents the mean and standard deviation for three replicates.

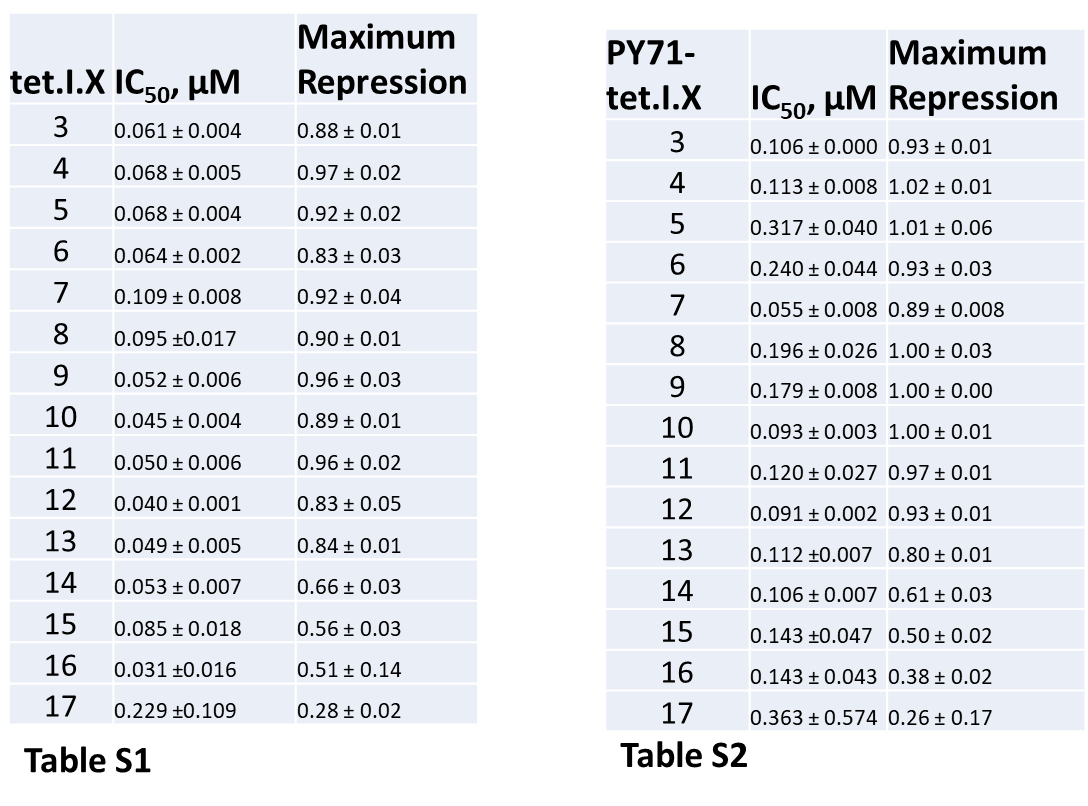
